# Pathways-wide genetic risks in Chlamydial infections overlap between tissue tropisms: A genome-wide association scan

**DOI:** 10.1101/205286

**Authors:** Chrissy h. Roberts, Sander Ouberg, Mark D. Preston, Henry J.C. de Vries, Martin J Holland, Servaas Morre

**Affiliations:** Department of Clinical Research, London School of Hygiene and Tropical Medicine, London, UK; Institute for Public Health Genomics, Department of Genetics and Cell Biology, School for Oncology and Developmental Biology (GROW), Faculty of Health, Medicine and Life Sciences, Maastricht University, 6200 MD Maastricht, The Netherlands; National Institute for Biological Standards and Controls, Potters Bar, UK; STI Outpatient Clinic, Department of Infectious Diseases, Public Health Service Amsterdam, Amsterdam, the Netherlands; Amsterdam Infection and Immunity Institute (AI&II), Academic Medical Centre, University of Amsterdam, Amsterdam, the Netherlands; Department of Dermatology, Academic Medical Center, University of Amsterdam, Amsterdam, the Netherlands; Laboratory of Immunogenetics, Department of Medical Microbiology and Infection Control, VU University Medical Center, 1081 HV Amsterdam, The Netherlands

## Abstract

*Chlamydia trachomatis* is the most commonly diagnosed bacterial sexually transmitted infection and can lead to tubal factor infertility, a disease characterised by fibrosis of the fallopian tubes. Genetic polymorphisms in molecular pathways involving G protein coupled receptor signalling, the Akt/PI3K cascade, the mitotic cell cycle and immune response have been identified in association with the development of trachomatous scarring, an ocular form of chlamydia-related fibrotic pathology. In this case control study, we performed genome-wide association and pathways based analysis in a sample of 71 Dutch women who attended an STI clinic who were seropositive for Chlamydia trachomatis antibodies and 169 high risk Dutch women who sought similar health services but who were seronegative. We identified two regions of within-gene SNP association with *Chlamydia trachomatis* serological response and found that GPCR signalling and cell cycle pathways were also associated with the trait. These pathway level associations appear to be common to immunological sequelae of Chlamydial infections in both ocular and urogenital tropisms. These pathways may be central mediators of human refractoriness to chlamydial diseases.

## INTRODUCTION

*Chlamydia trachomatis* (Ct) is diagnosed as the cause of around 106 million sexually transmitted infections (STIs) per annum [1] and has a significant impact on global health.

The primary pathology associated with Ct infection is an aberrant host immune response [2] which can lead to the progressive formation of scar tissues at and near the site of infection [3–5]. In the context of STIs this process can lead to infection related tubal factor infertility (TFI) and ectopic pregnancy [6]. Ct is also the leading infectious cause of blindness [7] and this form of disease (known as trachoma) is characterised by scar formation on the inner conjunctival surface of the upper eyelid, leading to lid deformation, corneal damage and visual impairment. Trachoma has historically been used as a platform for the study of chlamydial diseases and their pathology. This is largely due to the relative ease with which the conjunctiva can be studied compared to the fallopian tubes, but also because of the high prevalence of trachomatous scarring in sub-Saharan Africa [8, 9] compared to Ct pathology [10] in Europe and elsewhere.

Whilst serological evidence of historical Ct STIs can be detected in around a quarter of Northern European women without current Ct infections [10, 11] it can be very difficult to ascertain individuals for study who have laparoscopically confirmed cases of TFI with tubal scarring and concomitant seropositivity for historical Ct infections (Ct/TFI). In Europe the Ct/TFI prevalence in women is likely to be very low and in the Netherlands this was estimated at around 0.08% [10]. This makes the recruitment of participants to research studies focussed on urogenital chlamydial disease difficult. The collection of statistically powered case-control samples is extremely challenging but it is possible to leverage robust genetic association data from small samples by deploying statistical tests that not only collapse the highly dimensional genetic data to raise power [12], but also iterate over alternate scenarios to eliminate the burden of multiple testing [13]. We have used a pathways-based association test that not only corrects for the length of genes and pathways, but also utilises a process of permutation in which the genotypic associations of the observed traits were compared to a large number of simulated trait data sets. This process ensures that any pathways that were ultimately linked to the trait are not significant simply because of structure within the data, or because of limitations of the method, or because of the specific list of pathways that was chosen.

In our recent discovery phase genome-wide association study (GWAS) [14] we found that a number of genes and molecular pathways were associated with TS. That study applied both a classical single nucleotide polymorphism (SNP) association and systems based analysis [15–17] that focussed on the cumulative effects of multiple SNPs in the context of well-defined and curated molecular pathways of protein-protein interactions [18]. The trachoma GWAS identified genome-wide significant pathways that were associated with the protection and predisposition to TS. These results indicated a role for genes controlling the immune response, the mitotic cell cycle and numerous cell surface receptors and downstream signalling factors of the G-Protein Coupled Receptor family. The inference was that the key factors which protect the host from scarring were primarily innate barriers to initial infection and establishment of the intracellular niche, rather than direct protection from the scarring process itself. This interpretation raised a question about whether genetic protection might be so extensive in some people that they are innately refractory to becoming infected. In these people, we hypothesise that infection might never progress to the point that stimulates a lasting adaptive serological or cellular immune response.

The aim of this work was to assess the extent to which sexually transmitted Ct infections might share a common pathway of genetic predisposition with trachoma. By focussing on a trait of seropositivity for anti-Ct antibodies in high risk, presumably exposed Dutch women attending STI clinics, we investigated if pathways wide polymorphisms in the host are associated to primary events of infection. This work takes the form of a GWAS in a small case control sample in which we tested the association of 1345 multi-gene pathways, 590,811 directly genotyped SNPs and 8,852,410 imputed SNPs with serological status in 240 women attending a sexual health clinic in the Netherlands.

## METHODS

### Ethics statement

This study used archival material and linked data from clinical tests that had been irreversibly anonymised. The Medical Research Involving Human Subjects Act (WMO, Dutch Law) states that official approval of the study by the Medical Ethical Committee is not required for research using the anonymous human material collected for this study (MEC Letter reference: #10.17.0046). All subjects gave written informed consent for their specimens and data to be used in this way. All antecedent clinical studies were conducted in accordance with the tenets of the Declaration of Helsinki.

### Study population, sampling and ascertainment

244 Dutch Caucasian women were selected from a larger study consisting of Dutch Caucasian women visiting an outpatient sexual health clinic in Amsterdam, The Netherlands between July 2001 and December 2004. The women used a self-completion questionnaire to describe their symptoms at presentation, which varied from increased vaginal discharge, having bloody discharge during and/or after coitus, recent abdominal pain (not gastrointestinal or menses related) and/or dysuria.

### Genotyping and quality control

All specimens (n = 244) were genotyped at 713,599 SNPs using the Infinium OmniExpress-24 v1.2 Kit (Illumina). Basecalling was performed using Illuminus [19]. SNPs were retained in the analysis if (a) the call rate was ≥ 0.99, (b) the minor allele frequency was ≥ 0.99 and (c) Hardy-Weinberg equilibrium P value > 5 x 10^-8^. 590,811 directly genotyped SNPs were retained after QC.

PLINK and R were used to test for between-individual proportional IBS variance and to perform kinship estimation by identity by state/allele sharing estimation. Individuals were removed if they were identified as being outliers because they had: (a) > 10% missing genotype data. (b) Identity by descent (IBD) sharing with another individual of two alleles at all loci. (c) Identity by state with the fifth nearest neighbour with Z < -4 compared to the mean IBS of all possible pairs, or (d) unresolved gender mismatch between sex chromosome genotypes and clinical record. After these QC steps, 240 specimens were retained as one failed the pairwise IBD test and three failed the fifth nearest neighbour based outlier analysis.

### Statistical power

The R package “gap” [20] was used to estimate the power of the study to detect genome-wide significant ( < 1 x 10^-8^) associations of individual SNPs. At this level of significance, a study of 71 cases and 169 controls has 80% power to detect allele frequency odds ratios (OR) of 22.89, for minor allele frequency (MAF) in the control group of 0.1. The small sample size of this study meant that we could not differentiate between null results and false negatives. The false positive rate of SNP based genome-wide association tests was meanwhile controlled using the Bonferroni process. The false discovery rate of pathways analysis was controlled using a robust permutations process, which allows multiple testing at no increased cost to power[12, 13].

### Imputation

Imputation was carried out as described previously [21, 22] and included prephasing with Shapeit [23, 24] (informed by HapMap Phase II, build 37) and post-imputation filtering for high quality imputed SNPs with IMPUTE2 info score > 0.8 and/or MAF > 0.01 [14, 25]. Imputation was performed with IMPUTE2 [21, 22] using the worldwide 1000 Genomes phase I data set of 1092 reference specimens [26]. 20,514,944 SNPs were imputed and after filtering 9,443,221 SNPs were retained for association testing analysis.

### Tests of association

We modelled the clinical status (seropositive/seronegative) according to age using logistic regression. Residuals of the regression were taken forward as the serological status trait in the association test. Tests of association of SNPs with age corrected phenotypes were performed using EMMAX [27]. All tests of association were performed using a pairwise matrix of IBS kinship estimates which was generated from the SNP data. SNPNexus (http://snp-nexus.org) was used to annotate the SNPs with P < 1 * 10^-6^

### Pathways analyses

The R package SNPath (linchen.fhcrc.org/grass.html) was used to run the Assignment List GO AnnotaTOR (ALIGATOR) algorithm [12] on 1345 Reactome pathways [18]. In order to make the results directly comparable to those of the previous study [14], we chose to use the same pathways list that was used in that analysis (www.reactome.org, accessed April 2013). The P values from EMMAX tests of association were sampled to remove SNPs that were > 20 kb away from any gene in a list of ~17,000 genes that were consistently cross-referenced between Entrez and Ensembl and for which a HUGO gene name had been assigned (http://www.gettinggeneticsdone.com/2011_06_01_archive.html). ALIGATOR requires the user to specify a nominally significant threshold value and particularly when the sample size is low this choice can significantly affect the membership of the list of significant pathways. More stringent values for this threshold (i.e. 0.0001) will focus on SNPs most likely to be true positives (at the level of SNP) whilst more relaxed (i.e. 0.01) thresholds will capture more information from genes of small individual effect and may give a better indication of pathways involvement in diseases [12]. We chose to investigate the way in which threshold affected membership of the results table by running ALIGATOR at three thresholds of P_EMMAX_ (0.0001, 0.001, 0.01) and combining the results. We used 5000 permutations of the P_EMMAX_ values at each threshold. A list of all genes that appeared in any significant pathway at any threshold was generated and this list was functionally annotated using DAVID Bioinformatics Resources 6.7 [28] to identify enriched Reactome pathways and GO terms, whilst accounting for the significant gene content overlap in specific pathways.

### Serological and infection status determination

Peripheral venous blood was collected for the analysis of IgG antibodies against *C. trachomatis* (CT) using Chlamydia trachomatis-IgG-ELISA *plus* medac (Medac Diagnostika mbH, Hamburg, Germany). This test uses a synthetic peptide of a variable domain from an immune-dominant region of the major outer membrane protein (MOMP) and has been shown to perform with high specificity (~0.95-0.97) when evaluated against both Pgp3 and MIF [29]. A Ct IgG titre of ≥ 1:50 was considered positive. Samples with values within ± 10% of the 1:50 cutoff value were repeated and considered positive when the result was positive or still within the ± 10% range of the cutoff [30, 31]. The distribution of IgG titres in the positive population was as follows: Titre (T) = 1:50 [n = 9], T = 1:100 [n = 19], T = 1:200 [n=24], T = 1:400 [n = 9], T = 1:800 [n = 7] and T = 1:1600 [n=3]. In the negative population all specimens had a titre of T = 1:1 [n=169]. Tests for infection were performed on endo-cervical swabs using the COBAS AMPLICOR CT/NG Test, (Roche, Basel, Switzerland).

## RESULTS

There were 71 seropositive and 169 seronegative individuals (n = 240) in the analysis. All 71 seropositive cases were additionally infection positive, whilst all 169 seronegative cases were infection negative. Cases were significantly younger than controls in this sample (P = 0.007). The median age of cases was 23 (min 16, 1^st^ quartile 19.5, mean 22.6, 3^rd^ quartile 24.5, max 30). The median age of controls was 24 (min 15, 1^st^ quartile 21, mean 23.8, 3^rd^ quartile 27, max 32). The genome-wide inflation statistic was *λ* = 1.04 before correcting for age and *λ* = 1.02 after correction (Supplementary Figure 1).

In the genome-wide association study (GWAS), 46 single nucleotide polymorphisms (SNPs) with P_EMMAX_ < 1e-6 were identified (see Supplementary Table 1). These were located in five regions (A-E, see Table 1). Although none of the individual SNPs reached genome-wide significance (1e-8), regions B (Figure 1B) and C (Figure 1C) contained 33 and 12 SNPs respectively.

**Table 1.**
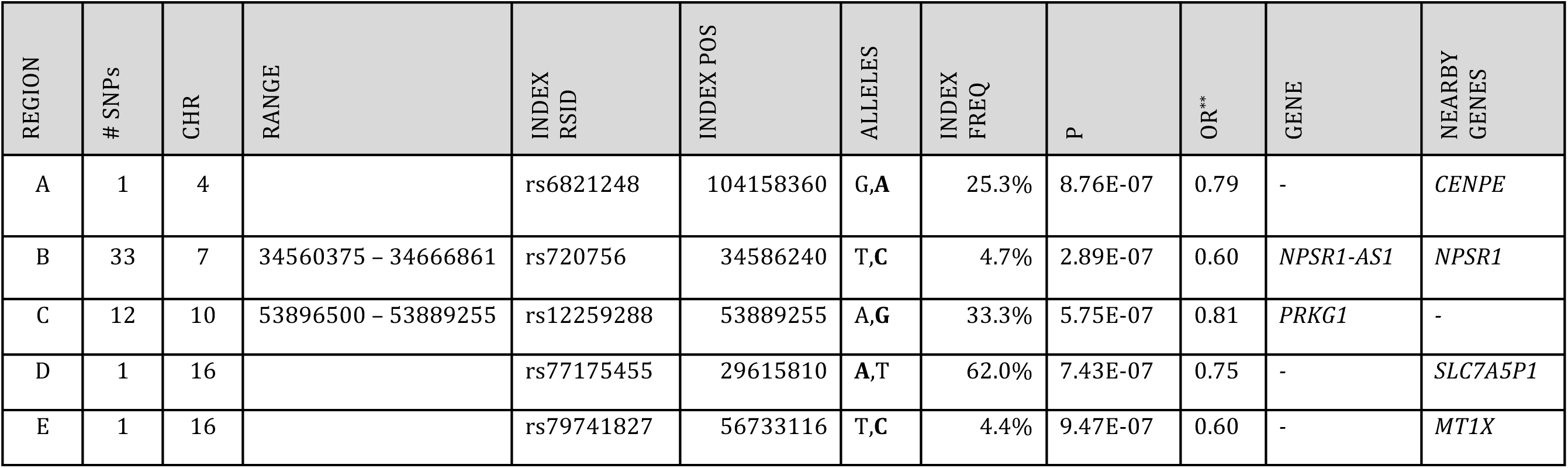
Index SNPs (P_emmax_ ≤ 1 x 10^-6^, at least one supporting SNP in LD with R^2^ > 0.6) for candidate associated regions. The index SNPs in the five genomic regions containing SNPs with P<1e-6. Indexes chosen by the lowest P_EMMAX_ in the region. SNP positions are relative to the Human Assembly 3GRCh37.p13 reference. All index SNPs were imputed.

| REGION | # SNPs | CHR | RANGE | INDEX<br>RSID | INDEX POS | ALLELES | INDEX<br>FREQ | P | OR** | GENE | NEARBY<br>GENES |
| --- | --- | --- | --- | --- | --- | --- | --- | --- | --- | --- | --- |
| A | 1 | 4 |  | rs6821248 | 104158360 | G,A | 25.3% | 8.76E-07 | 0.79 | - | *CENPE* |
| B | 33 | 7 | 34560375 – 34666861 | rs720756 | 34586240 | T,C | 4.7% | 2.89E-07 | 0.60 | *NPSR1-AS1* | *NPSR1* |
| C | 12 | 10 | 53896500 – 53889255 | rs12259288 | 53889255 | A,G | 33.3% | 5.75E-07 | 0.81 | *PRKG1* | - |
| D | 1 | 16 |  | rs77175455 | 29615810 | A,T | 62.0% | 7.43E-07 | 0.75 | - | *SLC7A5P1* |
| E | 1 | 16 |  | rs79741827 | 56733116 | T,C | 4.4% | 9.47E-07 | 0.60 | - | *MT1X* |

**Figure 1:**
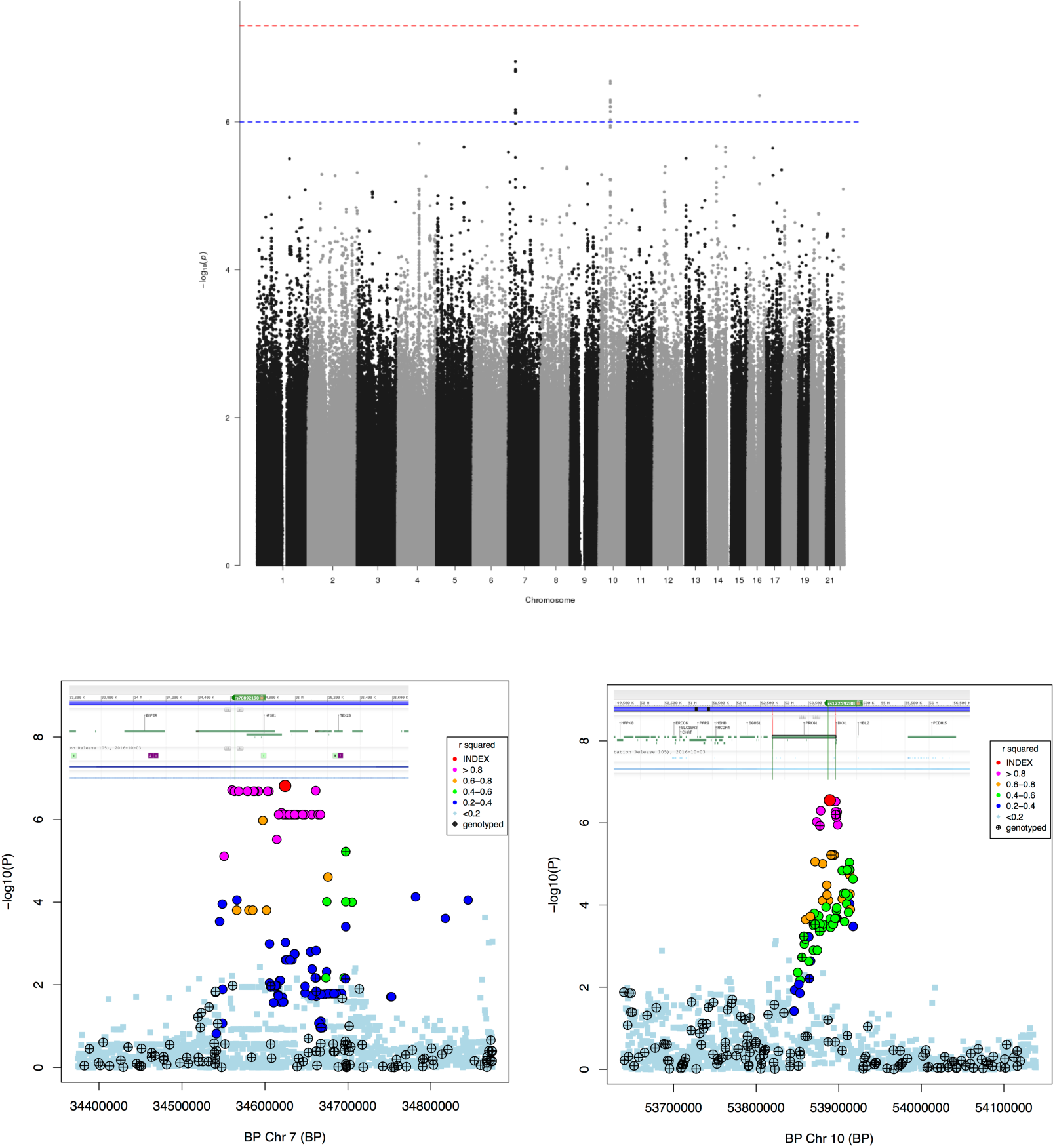
Results of GWAS Analysis for Chlamydia trachomatis seropositivity using EMMAX. (A) Manhattan plot showing index SNPs with P_EMMAX_ < 1 × 10-6. There were two regions of within-gene SNP association with Chlamydia trachomatis serological response. (B) The Chr 7: 34560375 - 34666861 region, within the non-protein coding NPSR1 Antisense RNA 1 (*NPSR1-AS1*) and immediately upstream of the G protein coupled receptor coding Neuropeptide S Receptor 1 (*NPSR1*). (C) The Chr 10: 53873323-56733116 region, within the gene encoding Protein Kinase, CGMP-Dependent, Type I (*PRKG1*), a mediator of the nitric oxide/cGMP signaling pathway, which has roles in immune function and GPCR signalling.

The 33 SNPs in region B lay within the non-protein coding NPSR Antisense RNA1 (*NPSR1-AS1*) and immediately upstream of the G protein coupled receptor coding Neuropeptide S Receptor 1 (*NPSR1*) (Figure 1B). The 12 SNPs in region C lie in the Protein Kinase, CGMP-Dependent, Type I (*PRKG1*) (Figure 1C), a mediator of the nitric oxide/cGMP signalling pathway, which has roles in immune function and in GPCR signalling.)

Sixty-six differently named Reactome pathways were identified as significantly enriched (P_perm_ < 0.05) in Chlamydial seropositivity. These pathways (Supplementary Table 2) are the combination of results from three ALIGATOR [12] pathway analyses with SNP cut-off thresholds of 0.01, 0.001 and 0.0001. The number of SNPs reaching each of the cutoff values were 93,106 (p<0.01), 9,998 (p<0.001) and 1168 (p<0.0001). The number of SNPs mapping to genes at each threshold were 64,397 (p<0.01), 6.274 (p<0.001) and 451 (p<0.0001). The number of significant pathways at each cut-off were 17 (p<0.01), 20 (p<0.001) and 33 (p<0.0001). In total, these pathways contained 2556 unique genes. Clustering the Reactome pathways by genic intersection demonstrated a large gene overlap (Figure 2).

**Figure 2:**
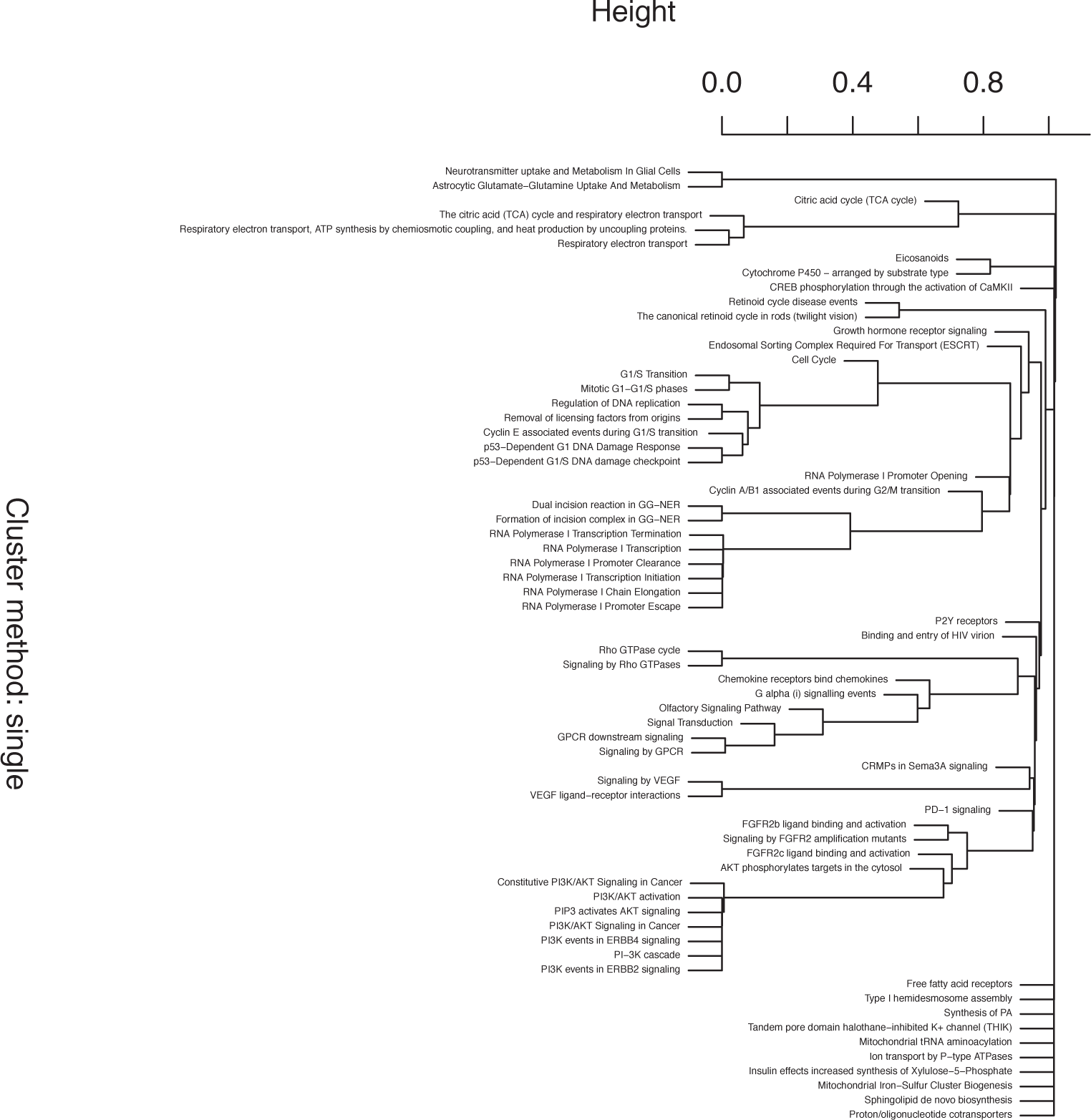
Gene content clustering of significant pathways in ALIGATOR analysis under three thresholds for nominal significance.

The Reactome “Signalling by GPCR” and “Cell Cycle, Mitotic” pathways contained 563 and 299 genes out of 2556 total genes (23.3% and 12.4%, respectively). Using the Gene Ontology (GO) database, over 10% of genes were assigned to the GO biological processes (GO-BP): “Cell surface receptor linked signal transduction”; “G-protein coupled receptor signalling pathway”; and “Sensory perception of smell”. This final process showing large genic overlap with the “Signalling by GPCR” Reactome pathway (Figure 2). The “Plasma membrane” (GO Cellular Component, GO-CC) and “Olfactory receptor activity” GO Molecular Function (Go-MF) are also enriched. The full list of significant pathways, with gene count and Benjamin Hochberg tests, can be found in Supplementary File 1.

## DISCUSSION

In this study we have identified two candidate regions of genetic association with anti-Ct serological status (P_EMMAX_ < 5 x 10^-6^) in Dutch women attending sexual health clinics for which there was at least one supporting SNP in LD with R^2^ > 0.6. One of these genomic regions of association was within the Protein Kinase, CGMP-Dependent, Type I gene (*PRKG1*) whilst another was within NPSR1 Antisense RNA 1 (*NPSR1-AS1*), which presumably has a role in post-transcriptional control of the nearby gene Neuropeptide S Receptor 1 (*NPSR1*).

The PRKG1 protein is a cytosolic cyclic GMP dependent protein kinase that has a role in signalling via the cRaf-MEK1/2-Erk-cMyc/cJUN axis. This may be important in Chlamydial disease as the trachoma GWAS [14] identified an association between pathways of signalling through Fibroblast Growth Factor receptors (FGFRs). We suggested that this system could be a crucial component in predisposition to trachomatous scarring. One pathway of signalling from FGFRs to the nucleus involves signal transduction through FRS2, Grb2/SOS and Ras, which then intersects with the cRaf-MEK1/2-Erk-cMyc/cJUN axis.

NPSR1 is a G-Protein Coupled Receptor that has been linked to Asthma susceptibility [32] and inflammatory bowel disease [33]. NPSR1 interacts with phospholipase C and like both *PRKG1* and *PREX2* (the best candidate associated gene from the trachoma GWAS [14]) is part of the G protein mediated cascade of intracellular signalling that centres around the PI3K/Akt pathway of cell cycle regulation. Kechagia and colleagues showed that trachoma fibroblasts could promote Akt phosphorylation in macrophages in an IL-6-dependent manner [34]. They went on to suggest that increased IL-6 secretion in trachoma may sustain macrophage activation and survival, with the knock-on effect that this could enhance the contractile activity of the fibroblasts and lead to more severe scarring. Whether the role for the PI3K/Akt pathway in trachoma is to protect against primary infection, or against progression to scarring, or both things, is a matter that requires further study.

This study has some significant limitations. The sample used in this study was very small and the SNP based analyses presented in this paper are therefore critically underpowered. Whilst we are confident that the positive signals from the *PRKG1* and *NPSR1-AS1* region are interesting enough to warrant further study, we recognize that the false negative rate is very high. To leverage additional power from the small sample, we focused on pathways-wide analysis that was similar to the one used previously [14]. We determined that the ALIGATOR method was somewhat unstable with regards to the specific pathways that were highlighted at different thresholds of nominal significance (Supplementary Table 2). This is no doubt because more permissive nominal cut-off values (i.e. p<0.01) lead to a more sensitive analysis that identifies more pathways but lacks the more granular specificity of more conservative thresholds (i.e. p<0.0001). Collating the results of three different runs of the ALIGATOR analysis allowed us to average across the various sources of error (false negatives vs false positives) whilst identify pathways.

When we assessed the data with functional analysis using DAVID [28], the results pointed towards the importance of cell cycle and G protein coupled receptor mediated signalling (Supplementary File 1). By testing for a limited number of pathways whilst utilizing permutation based estimators of significance, we have been able to robustly detect signals of pathways wide association with chlamydial seropositivity that are almost identical to the findings of the previous study in trachoma.

We identified potential roles for pathways relating to PI3K/Akt signalling, FGFRs and control of the cell cycle. G-PCRs and their signalling pathways were previously highlighted in trachoma cases [14, 35] and also in *Chlamydia muridarum* infection in the BXD recombinant inbred mouse [36]. Neural growth factor receptor (NGFR) pathways were also strongly associated with serostatus in the Dutch population. Derrick and colleagues [37] previously showed that numerous gene and pathway targets of trachoma associated microRNA moieties were neuronal related.

We also detected a significant role for olfactory receptor signalling cascades in this study (Supplementary File 1). This could be accounted for by the substantial overlaps between the genes involved in olfactory signalling and the G protein/PI3K signalling system. The olfactory system has however been independently linked to *C. pneumoniae* infections and subsequent fibrillogenic Alzheimer’s like plaque formation in the olfactory centres of mouse brains [38]. Given that such plaques involve collagen restructuring and might be considered akin to the scars that succeed human Ct infections, it seems reasonable to infer that the various *Chlamydiaceae* can utilise a variety of different, but related, cell surface receptors to gain access to the host cell at different body sites. These diverse receptors may well share a common downstream signalling process via G proteins and particularly via the PI3K pathway. Collectively these data appear to confirm that G protein coupled receptor signalling pathways and the cell cycle are associated with chlamydial infections in a way that transcends tissue tropisms and which might be utilised by many or all chlamydial species in their animal hosts. In the absence of an appropriate replication panel for the more granular analyses of the SNP based association tests, we infer more meaning from the functional overlap of the pathways identified in this study and the previous GWAS in trachoma. We interpret our data in the broadest sense, which is that the key pathways identified between the studies have substantial functional and gene content overlap. A cross-study comparison of the 'p values’ for any specific pathway is not however an appropriate way to interpret this overlap, largely because the high false negative risk leads to a certain instability in the association test results. With this small study, we are unfortunately not able to quantify the convergence of the results between the studies.

Another limitation of the study that we highlight is that trait of sero-positivity was measured using a single antigen (MOMP) test. It was not possible to prove unequivocally that all the women who were included in the study as controls had been exposed to *C. trachomatis* infections, nor that co-infection with other micro-organisms was not a complicating factor in the clinical presentation of our cases, but this study was carried out in the context of a population of women seeking test and treatment, who were thought to be at very high risk for exposure to *C. trachomatis* STI. Our control group had a uniform antibody titre of 1:1, whilst the majority of positives had high titres (mode 1:200).

We have selected statistical analysis methods for the pathways analysis that allow for a high multiplicity of testing, whilst not imposing the same burden of signal loss that are associated with more rudimentary corrections such as Bonferroni’s method. They also collapse highly dimensional data, leading to a better powered 'burden' test. In the light of the robust permutations based analysis we used the remarkable similarities between our findings and those of the previous trachoma GWAS, it is extremely unlikely that these associations have been identified in both the trachoma and STI tropisms either by chance or because of systematic errors in our analysis approach.

By focusing on a trait that is generally accepted to indicate lifetime responses to infectious challenge, we believe that we have identified pathways-wide genotypes that appear to associate with protection from infection in some women. This protection could take the form of a partial or complete refractoriness to primary cellular infection, or a rapid, intense and highly effective innate immune response.

### Conclusions

Between the tissue tropisms of the eye and the genital tract, the pathways-wide genetic risk and protection patterns overlap to a substantial degree. This supports a model under which common pathways of intracellular signalling might be utilised by all *Chlamydiaceae,* even though the specific receptors that mediate introgression in to the cell might be highly variable.

## Acknowledgements

We are very grateful to the participants of this study.

## Author Contributions

S.M., S.O., M.J.H & C.h.R.designed the study. S.O., S.M. & HJCdV performed the clinical collections and laboratory tests. M.D.P., M.J.H. & C.h.R. performed the computational work and statistical analyses. All authors wrote, reviewed and approved the manuscript.

